# Sex-dependent and region-specific morphological adaptations in astrocytes following early-life stress

**DOI:** 10.64898/2026.11.05.764658

**Authors:** Camila Cabarcas Arrieta, Anthony Bosson, Mathias Guayasamin, Kathleen Ngo, Sarah Peyrard, Ciaran Murphy-Royal

## Abstract

Astrocytes are structurally heterogeneous, with cell morphology providing a valuable readout of experience-dependent plasticity. Despite this, single-cell reconstruction including complete cellular territories remains challenging to compare across brain regions and experimental groups. We used unilateral retro-orbital delivery of PHP.eB-GfaABC1D-eGFP as a practical method for sparse astrocyte labeling and applied it to evaluate regional, sex, and early-life-stress comparisons in the mouse amygdala. Systemic virus delivery to broadly target astrocytes across the brain produced eGFP-positive cells throughout cortex, hippocampus, hypothalamus, and amygdala with spatially isolated cells suitable for intensity projection, three-dimensional surface reconstruction, filament tracing, and Sholl analysis. In naive females, we report relatively homogenous morphological features between lateral amygdala and basal amygdala. In naive males, basal amygdala astrocytes had greater volume and a lower surface-area-to-volume ratio, while surface area, soma diameter, territory, and arborization measures did not differ significantly. Interestingly, early-life stress produced sex-specific morphological responses in the basal amygdala. Females showed increased perimeter, area, and Feret diameter without significant changes in three-dimensional volume or resolved filament measures. Males instead showed reduced surface area and volume, with no significant changes in projected territorial dimensions, soma diameter or resolved filament measures. S100β-positive cell density was unchanged in the displayed regional and stress comparisons. These findings establish systemic sparse eGFP labeling as an efficient route to single-cell astrocyte morphometry and reveal a sex-dependent and region-specific dissociation between territorial expansion and three-dimensional volumetric contraction after early-life stress.

**MAIN POINTS:**

- A single retro-orbital administration of AAV.PHP.eB-GfaABC1D-eGFP labels spatially isolated astrocytes across multiple brain regions and supports surface, filament, and Sholl analysis.
- Under naive conditions, male basal amygdala astrocytes have greater reconstructed volume and lower surface-area-to-volume ratio compared to lateral amygdala astrocytes, whereas the corresponding female regional comparisons are not different.
- Early-life stress is associated with sex-dependent BLA remodeling. Females exhibit greater territory size, whereas males show reduced three-dimensional surface area and volume.

## INTRODUCTION

Astrocytes display complex morphologies following a general structural motif composed of a soma, major branches, progressively finer branchlets and peripheral processes, which comprises of at least a single endfoot wrapping vasculature [1–3]. This morphological complexity and high degree of compartmentalization enables discrete interactions between astrocytes and synapses, blood vessels and the extracellular space across the central nervous system regions [4,5]. Morphology is therefore a useful readout of local specialization and experience-dependent plasticity, but it is technically difficult to distinguish individual territories, where astrocytes interface with each other at the distal processes [3,6]. GFAP immunostaining identifies major processes and reactive changes, yet it labels only a fraction of the volume of many protoplasmic astrocytes [7,8]. Methods that fill the cytoplasm more completely improve morphometry but often require transgenic lines, intracellular filling or local injections targeted separately to each region [8].

Here we used a viral construct, shown to efficiently cross the blood-brain barrier after systemic delivery in mice [9–11], to drive eGFP expression in astrocytes by employing a GfaABC1D promoter [12–15]. We asked whether this approach could achieve sufficient anatomical targeting and single-cell resolution to compare astrocyte morphology between neighboring amygdala subdivisions and to detect sex-dependent remodeling after early-life stress. Previous work has shown that stress can reshape astrocyte structure and that structural responses differ across stress paradigms and sexes [16–20], but detailed comparison of subregions under unstressed conditions remains limited. The lateral amygdala (LA) and basal amygdala (BA) are neighboring components of the basolateral amygdala complex with distinct circuit organization and contributions to emotional learning [21–24]. Astrocytes within these regions may therefore face different functional demands.

As a first step to address potential heterogeneity in astrocytes within the BLA, we mapped reporter distribution across the brain and established a reconstruction workflow combining projected territorial measurements, three-dimensional surface models, filament models, and Sholl analysis. We then compared LA and BA astrocytes separately in naïve (non-stressed) females and males. Finally, we examined BLA astrocytes after early-life stress, again separating the analyses by sex. The resulting data identify a male-specific regional difference under naive conditions and distinct female and male structural responses after early-life stress.

## MATERIALS AND METHODS

### Animals and experimental design

Male and female C57BL/6J mice (The Jackson Laboratory, stock no. 000664) were housed on a 12-h light/dark cycle (lights on at 06:30) with food and water available ad libitum. All procedures were performed in accordance with Canadian Council on Animal Care guidelines and were approved by the Institutional Committee for the Protection of Animals at the Centre hospitalier de l’Université de Montréal under protocol CM20031CMRs. Experimental groups comprised naive females, naive males, early-life-stressed females and early-life-stressed males.

### Early-life stress

Early-life stress was performed following the paradigm introduced by [25] that we previously validated to affect amygdala-dependent behaviour [16]. C57BL/6J pups were separated from their dams for 4 h per day (ZT2-ZT6) on seven consecutive days from P10 to P17. During each separation, dams and pups were placed in separate, prepared cages, each containing 33% of the home cage’s original bedding. On P10, the litter’s bedding allotment was weighed and divided approximately equally among the home cage, pup-separation cage and dam-separation cage. Separated pups and dams had *ad libitum* access to food and water, and pup cages were placed on a water-heating pad maintained at 34°C. After the final separation on P17, the original bedding amount was restored in the home cage, and pups were rehoused with their dams until weaning at P21.

### Viral vector and retro-orbital administration

Retro-orbital delivery was performed following established principles described by [26]. On postnatal day 30, mice were anesthetized by inhalation with isoflurane (2%, v/v) until loss of the toe-pinch reflex. A drop of topical local anesthetic (0.5% proparacaine hydrochloride ophthalmic solution, Alcon Laboratories) was applied on the designated eye and left in place for approximatively one minute before proceeding with the injection. The syringe was inserted at a 30-degree angle, bevel facing down, into the medial canthus until reaching the retroorbital sinus at the base of the eye. Mice received 50 µl of viral vector AAV.PHP.eB-GfaABC1D-eGFP (1 × 10^11-12^ GC/mL) produced by CERVO Canadian Optogenetics and Vectorology Foundry Core Facility (RRID:SCR_016477). Afterward, rodents were single housed until anesthesia wore off.

### Tissue preparation and immunohistochemistry

Mice were perfused at P51, 21 days after vector delivery. Animals were anesthetized in an enclosed chamber with isoflurane (5% for induction and 3-4% for maintenance, v/v) until loss of the toe-pinch reflex. Following transcardial perfusion, brains were fixed in 4% paraformaldehyde for 24 hours at 4°C and cryoprotected in 30% sucrose for at least 2 days. Brains were embedded in optimal cutting temperature compound, flash-frozen in 2-methylbutane maintained at -50°C and stored at -80°C until cryosectioning. Coronal sections (30 μm) were cut at -20°C on a Leica CM3050S cryostat. They were subsequentially placed in wells containing 800 µl of 1X phosphate-buffered saline (PBS) solution at 4°C until immunohistochemistry.

Free-floating sections were washed three times for 15 min in 1X PBS to remove cryoprotectant, then, blocked and permeabilized for 1 h in a 1X PBS solution containing 3% bovine serum albumin and 0.5% 10X Triton. Sections were incubated for 24 h at 4°C in a solution of primary antibodies. For slices destined for morphological assessment, the antibodies used were rabbit anti-eGFP (1:500, Thermo Fisher Scientific, G1362) and chicken anti-GFAP (1:1000, Thermo Fisher Scientific, PA1-10004). Slices intended for cell density analysis were incubated in a solution containing rabbit anti-S100β (1:1000, Abcam, AB5264). After three 15-min PBS washes, sections were incubated for 1 h at room temperature, protected from light, in a PBS solution with DAPI (1:10,000) and species-appropriate fluorophore-conjugated secondary antibodies (Morphometry slices: goat anti-rabbit Alexa Fluor 488 (1:1000, Jackson ImmunoResearch Laboratories, 111-545-144) and goat anti-chicken Alexa Fluor 568 (1:1000, Thermo Fisher Scientific, A11041; Cell density slices: goat anti-rabbit Alexa Fluor 488 (1:1000, Jackson ImmunoResearch Laboratories, 111-545-144 )). Sections were washed three further times for 15 min in PBS and mounted on Fisherbrand microscope slides using ProLong Glass Antifade mountant (Thermo Fisher Scientific, P36982).

### Imaging and anatomical sampling

Images for single-cell morphometry analysis were acquired using Leica Microsystems Stellaris 8 inverted confocal microscope. It was equipped with an HC PL Apo CS2 63×/1.40 Oil objective and a TD 488/561/638 spectral separator. Excitation was performed with 405, 488, 561 and 638 nm lasers and HyD detectors with 493-532 nm, 415-483 nm and 581-726 nm spectral windows were ensuring detection. The pinhole was set to 1 Airy Unit (AU). Acquisition was sequentially conducted with the excitation and detection of the 488 nm channel (eGFP) first and then, both 405 nm (DAPI) and 561 nm (GFAP) channels. Images were obtained in frame mode, with a 1× zoom and a scan speed of 1,57 µs/pixel in unidirectional mode. For the 488 nm channel, line accumulation was adjusted at 2, whereas frame accumulation, line averaging and frame averaging were set to 1. For the 405 and 561 nm channel, all parameters were maintained at 1. Image resolution was 1024 × 1024 pixels with a depth of 16 bits. Pixel size was 0,18 µm × 0,18 µm. Z-stacks contained between 72 to 121 planes which were each separated by 0,30 µm. Each image was saved using the LAS X software (version 4.6.1.27508).

Images used for cell density analysis were acquired using a Zeiss Observer Z1 spinning disk confocal microscope with a 20X 1.8 NA objective. Images were acquired as tiled z-stacks, each tile consisting of 27 stacks acquired at 0,68 μm step, for a total depth of 17,68 μm. DAPI fluorescence was acquired using 405 nm channel at 3% laser power with an exposure time of 200 ms, while S100β fluorescence was acquired through the 488 nm channel at 9% laser power with an exposure time of 800ms. Imaging parameters were consistent across all samples.

Images of coronal brain sections were acquired using an Aperio VERSA fluorescence slide scanning microscope (Leica Biosystems) equipped with a Leica HC PLAN APO 20x dry objective (0.75 NA). Images of multiple mouse brain coronal sections on a single slide were captured in fluorescence mode at a native pixel resolution of 0.32 μm/pixel. To account for tissue topography, autofocusing was performed using a 7-plane focal map with a 0.7 μm step size. Imaging and stitching parameters were kept identical across all samples.

### Cell density and selection for reconstruction

S100β-positive cell density was measured in the LA and BA of naive mice, as well as in the BLA of and early-life-stressed animals. A maximum intensity projection was used to create a two-dimensional image. A region of interest (ROI) was hand-drawn for both regions, using the external and amygdala capsule to distinguish them. ImageJ (2.16.0/1.54p) was utilized to count cell numbers in each region with the Analyze particles tool. Detections were then manually reviewed to remove false-positive and recover false-negative profiles. Obtained number of S100β-positive cells were normalized by dividing them by the volume, which itself was a multiplication of the ROI, the number of stacks and the size interval in between stacks. Number of stacks varied between 15 and 32. Results are expressed in number of cells per cubic millimeter.

### Morphological reconstruction

Image stacks were processed in the software IMARIS (version 10.2). Surface area, volume and surface area to volume (SA/V) were determined by reconstructing a Surface model of astrocyte volume using Surface rendering. Surface detail level was set to 0,118 µm. Absolute intensity threshold varied between 101 and 209. Additionally, seed point diameter was 0,3 µm. Soma diameter was measured by the ruler tool in the software, which was necessary for the creation of a filament model using the Filament Tracer. This 3D reconstruction of astrocyte processes was then used to perform a Sholl analysis and to calculate total process length and number of branch points. Three-dimensional Sholl analysis quantified intersections as a function of radial distance from the soma, and the maximum number of intersections was extracted for group comparison. Seed point diameter was set to 0,3 µm and starting points threshold ranged from 250 to 395. Astrocytes overlapping other eGFP-positive cells without clear demarcation were not included. Endfeet were manually excluded from analysis, as the reconstruction models could not capture the specificity of the structure.

Images used for the three-dimensional reconstructions also served for the two-dimensional assessments. A z-stack was created using maximum intensity projection. Single cell outline was hand-drawn and the measurement tool in ImageJ calculated values for the perimeter, the area and the Feret diameter.

### Statistical analysis

Baseline regional comparisons were performed separately in females and males. Stress comparisons were performed within female BLA and male BLA, followed by a joint model testing the sex-by-stress interaction for each prespecified end-point. Detailed statistical tests are included in the figure legends. Bars in Figures 2-5 show group means with error bars, and individual observations are superimposed. Heatmaps summarize z-scored group means across astrocyte density, projected territory, surface, soma, filament and Sholl-derived endpoints. For groups respecting normality, two-tailed Student’s t-test were executed when group variances were assumed equal, otherwise, Welch’s t-test was selected. Data with non-Gaussian distribution was analyzed with Mann Whitney test. All data is presented as mean ± SEM, unless expressed otherwise. Statistical analyses were executed using GraphPad Prism (version 11.1.0). Statistical significance is indicated the following way: * for p < 0.05, ** for p < 0.01 and *** for p < 0.001.

## RESULTS

### Systemic delivery provides distributed sparse labeling for single cell morphometry

To establish whether systemic virus delivery could support detailed astrocyte morphometry, mice received a single retro-orbital injection of AAV.PHP.eB-GfaABC1D-eGFP at postnatal day 30 and were perfused three weeks later, followed by immunohistochemical processing and imaging (**Fig. 1A,B**). Whole-section epifluorescence imaging revealed reporter-positive cells across the forebrain, including cortex, hippocampal CA2, hippocampal stratum radiatum, lateral hypothalamus, arcuate nucleus and basolateral amygdala (**Fig. 1C**). Expression was regionally heterogeneous, but each sampled structure contained spatially isolated cells with a soma and complex peripheral eGFP signal along astrocyte processes (**Fig. 1C, insets 1-6**). Subsequent imaging, using confocal microscopy, showed the extent of reporter-filled astrocyte profiles in relation to conventional GFAP labeling (**Fig. 1D**). Finer processes were better observed with the eGFP labelling as opposed to the GFAP staining, which reveals an incomplete structure [27]. Indeed, co-staining with GFAP, enabled single cell morphological reconstruction with compartment specific identification of endfoot processes (**Fig. 1E**). These data support the use of a single systemic administration of astrocyte-targeting virus to provide morphological access to multiple brain regions, yielding isolated cells suitable for single-cell analysis.

**Figure 1.**
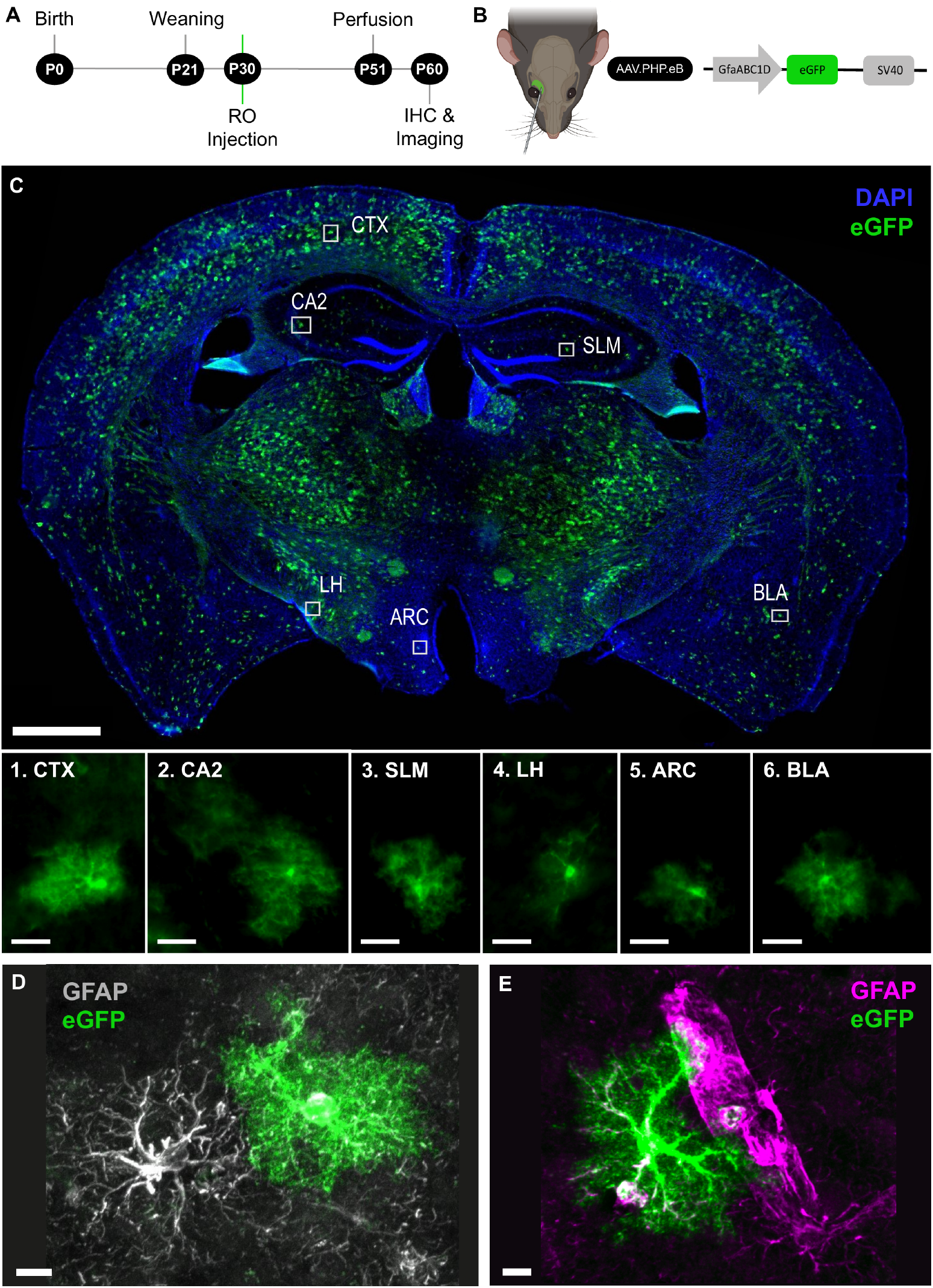
Systemic AAV.PHP.eB-GfaABC1D-eGFP delivery labels spatially isolated astrocyte like cells across the brain. **(A)** Experimental timeline. Mice were weaned at P21, received retro-orbital vector at P30, were perfused at P51 and underwent immunohistochemistry and imaging by P60. **(B)** ERetro-orbital delivery and vector organization showing GfaABC1D-driven eGFP and an SV40 regulatory sequence. **(C)** ECoronal section showing DAPI and eGFP signal, with cortex (CTX), hippocampal CA2, stratum lacunosum-moleculare (SLM), lateral hypothalamus (LH), arcuate nucleus (ARC) and basolateral amygdala (BLA) indicated. Scale bar: 900 µm. Boxes identify representative reporter-positive cells enlarged in insets 1-6. Scale bars: 20 µm. **(D**,**E)** Representative eGFP-positive astrocyte-like profiles shown in GFAP-labeled fields. Scale bars: 10 µm.

### Astrocytes in lateral and basal amygdala have similar morphologies in female mice

We focused on the amygdala, a brain region which we, and others, have shown astrocytes play central roles in synaptic and behavioural output [13–16,19,28]. We compared lateral amygdala (LA) and basal amygdala (BA) astrocytes in naive females (**Fig. 2A,B**). Using a pan-astrocytic marker S100β we found no differences in cell density between regions (**Fig. 2C-D**). Next, single-cell z-stack were projected at maximum intensity to obtain a two-dimensional image for more typical assessment of astrocyte perimeter (the outline of labelled cell), area (the space occupied by the surface inside the traced perimeter) and Feret diameter (the longest distance between two opposing points on the cell), reflective of cell elongation (**Fig 2E**). Using two-dimensional analysis, we report that astrocytes in the LA and BA present similar morphological characteristics (**Fig. 2F-I**). Next, we carried out three-dimensional morphometry, analysing surface area, volume, surface-area-to-volume ratio, and soma diameter (**Fig.2J**). Despite a trend in volume, we observed no difference in astrocytes originating from either region (**Fig. 2K-N**). Using a filament model, total process length, branch-point number, and maximum Sholl intersections were likewise unchanged (**Fig. 2O-S**). The Sholl profiles overlapped broadly across radial distances despite a descriptive difference in their plotted maxima (**Fig. 2T**), which could relate to trends observed in three-dimensional volume. Endfeet structures were manually removed from created filament and surface models, as we could not generate an accurate representation of the subcellular compartment. The integrated heatmap summarizes the data, with an absence of robust differences across the morphological features (**Fig. 2U**). Overall, the present dataset suggests homogenous astrocyte density and morphology, using both two- and three-dimensional analyses, between female LA and BA astrocytes.

**Figure 2.**
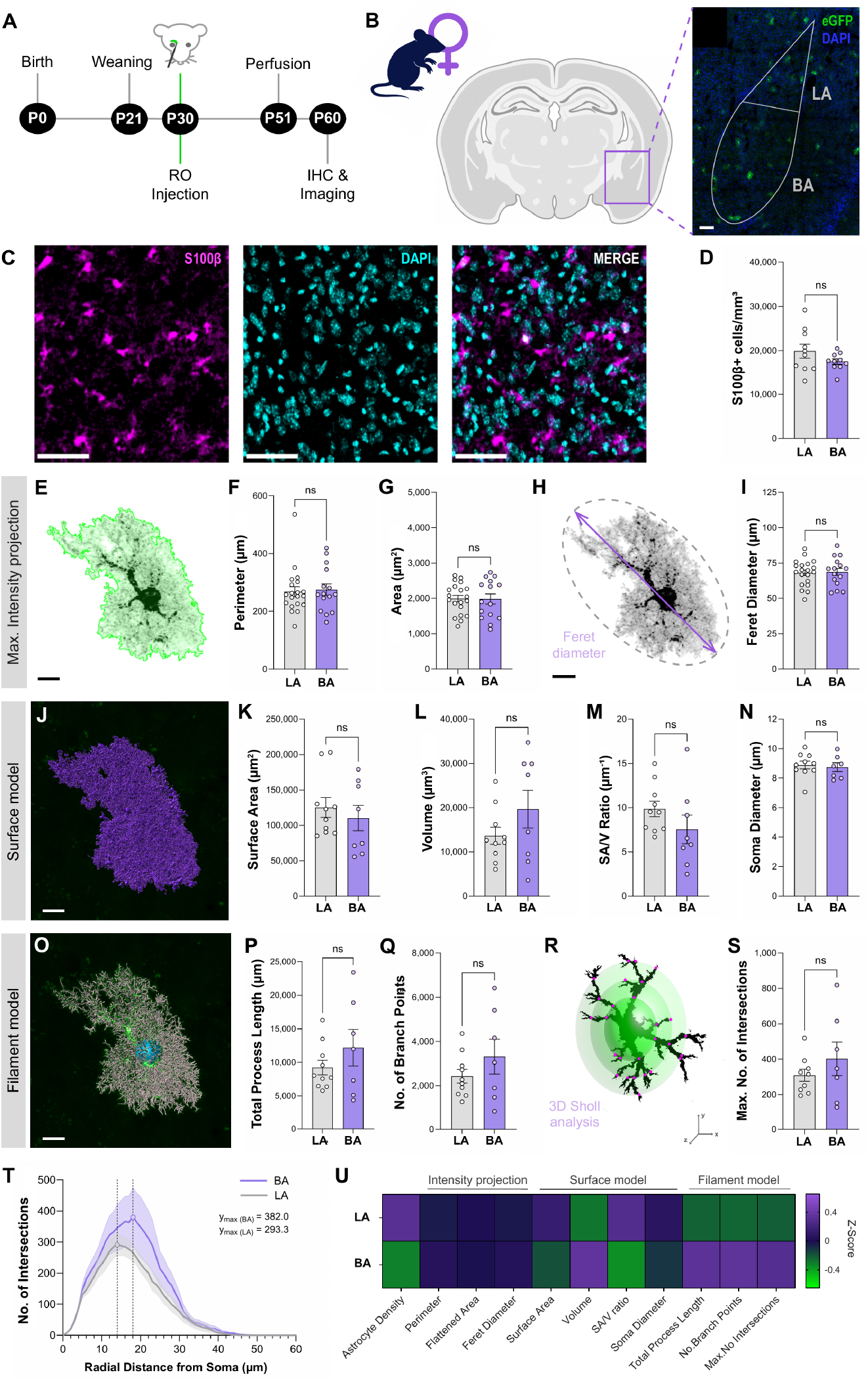
Naive female LA and BA astrocytes have similar morphometric measurements. **(A)** Experimental timeline. **(B)** Anatomical sampling of lateral amygdala (LA) and basal amygdala (BA), with representative eGFP and DAPI signal. Scale bar: 100 μm. **(C)** S100β-positive cells, DAPI and merged images taken within the BLA. Scale bars: 50 μm. **(D)** S100β-positive cell density in both regions (Welch’s t test: LA = 19,900 S100β+ cells/mm^3^ (N = 7, n = 10 slices), BA = 17,540 S100β+ cells/mm^3^ (N = 7, n = 10 slices), p = 0.1861). **(E)** Representative maximum intensity projection used for perimeter, area, Feret diameter measurements. Scale bar: 10 μm. (F, G) Perimeter (Mann-Whitney test: LA = 255.7 μm (N = 5, n = 20 cells), BA = 258.7 μm (N = 5, n = 15 cells), p = 0.8307) and area (unpaired t test: LA = 1,991 μm^2^ (N = 5, n = 20 cells), BA = 1,982 μm^2^ (N = 5, n = 15 cells), p = 0.9573). **(H, I)** Feret diameter definition and quantification (unpaired t test: LA = 68.04 μm (N = 5, n = 20 cells), BA = 68.70 μm (N = 5, n = 15 cells), p = 0.8405). Scale bar: 10 μm. **(J)** Representative three-dimensional surface model created to calculate surface area, volume, surface-area-to-volume ratio. Scale bar: 10 μm. **(K-N)** Surface area (Mann-Whitney test: LA = 115,500 μm^2^ (N = 5, n = 10 cells), BA = 98,650 μm (N = 5, n = 8 cells), p = 0.3722), volume (unpaired t test: LA = 13,645 μm^3^ (N = 5, n = 10 cells), BA = 19,664 μm^3^ (N = 5, n = 8 cells), p = 0.1874), surface-area-to-volume ratio (unpaired t test: LA = 9.856 μm^-1^ (N = 5, n = 10 cells), BA = 7.531 μm^-1^ (N = 5, n = 8 cells), p = 0.1973) and soma diameter (Mann-Whitney test: LA = 8.945 μm (N = 5, n = 10 cells), BA = 8.810 μm (N = 5, n = 7 cells), p = 0.5362). **(O)** Representative filament model used to obtain total process length, number of branch point and three-dimensional Sholl analysis. Scale bar: 10 μm. **(P, Q)** Total process length (Mann-Whitney test: LA = 7,881 μm (N = 5, n = 10 cells), BA = 12,000 μm (N = 4, n = 7 cells), p = 0.6691) and branch-point number (Mann-Whitney test: LA = 2,185 (N = 5, n = 10 cells), BA = 3,052 (N = 4, n = 7 cells), p = 0.6009). **(R)** Sholl analysis schematic. **(S)** Maximum number of intersections (Welch’s t test: LA = 308.7 (N = 4, n = 9 cells), BA = 401.9 (N = 5, n = 7 cells), p = 0.3823). **(T)** Sholl-intersection profile across radial distance. **(U)** Summary heatmap of z-scored group means across cell density and the displayed morphological endpoints. Bars show group means with SEM, and points show individual observations. All displayed comparisons are not significant. ns, not significant.

### Astrocytes exhibit regional differences in the amygdala of male mice

Our subsequent analysis focused on astrocytes from the brains of male mice. The comparison of LA to BA astrocytes in naive males revealed selective regional difference (**Fig. 3A,B**). As in female mice, we observed no change in astrocyte density across the BLA complex, evidenced by similar S100β-positive cell densities across the LA and BA (**Fig. 3C, D**). Indeed, single cell analysis revealed that individual astrocyte perimeter, two-dimensional area, and Feret diameter (elongation) were comparable between brain regions (**Fig. 3E-I**). While three-dimensional surface area was similar across regions (**Fig. 3J, K**), BA astrocytes had greater three-dimensional volume than LA astrocytes (**Fig. 3L**). This greater volume, associated with a similar area, resulted in a significantly lower surface-area-to-volume ratio (**Fig. 3M**). These findings suggest regional differences in the relationship between surface area and volume. Soma diameter, total process length, branch-point number and maximum Sholl intersections did not differ significantly between regions (**Fig. 3N-S**), with Sholl profiles broadly overlapping (**Fig. 3T**). The integrated heatmap of our findings highlights that the male regional profile is driven principally by the opposing volume and surface-area-to-volume signals rather than a generalized shift across all morphological measures (**Fig. 3U**). Together, these data indicate greater three-dimensional volume in male BA astrocytes without a broad increase in resolved arbor complexity.

**Figure 3.**
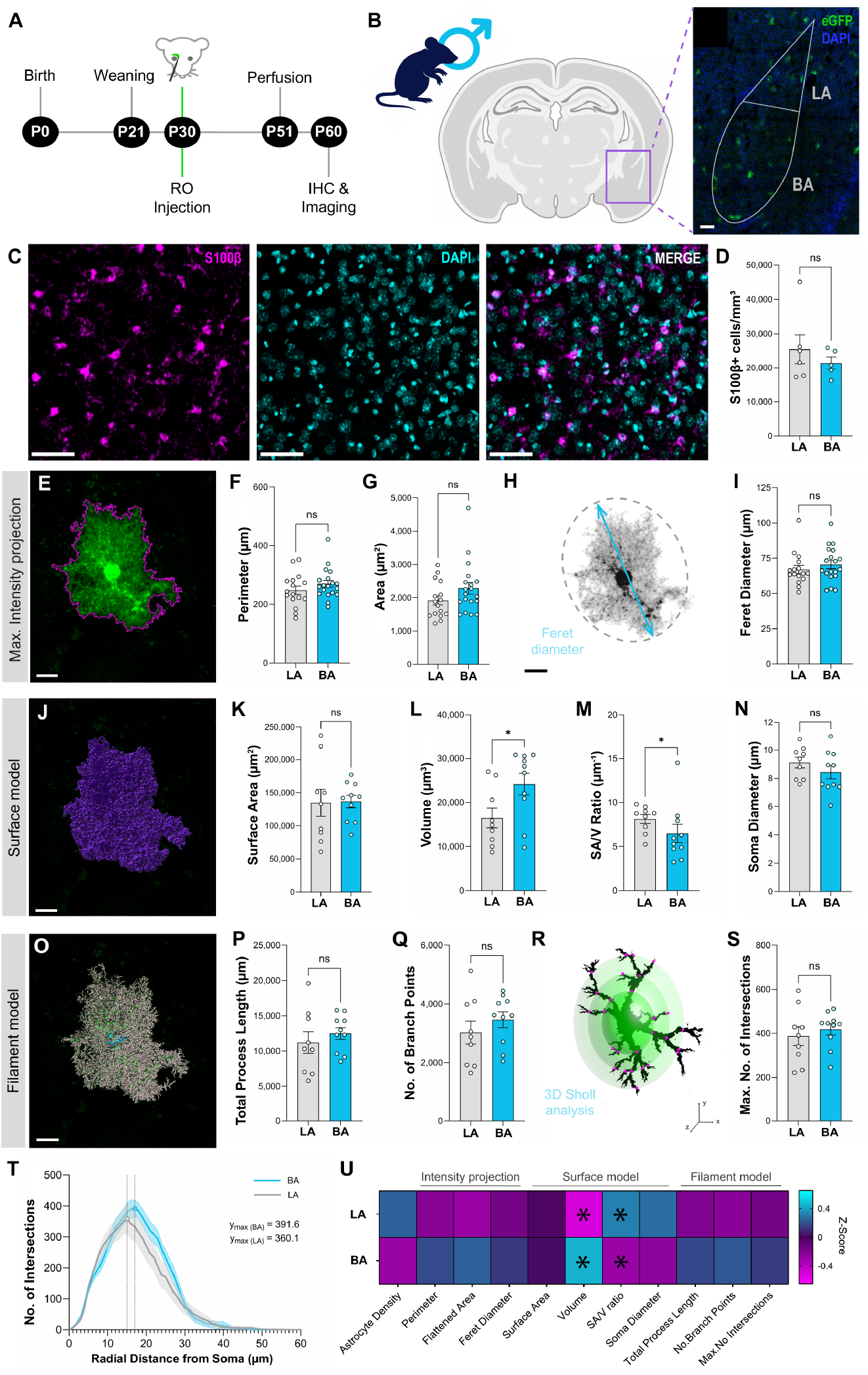
Naive male BA astrocytes have greater volume and lower surface area to volume ratio. **(A)** Experimental timeline. **(B)** Anatomical sampling of LA and BA, with representative eGFP and DAPI signal. Scale bar: 100 μm. **(C)** S100β, DAPI and merged images. Scale bars: 50 μm **(D)** S100β-positive cell density (Mann-Whitney test: LA = 23,800 S100β+ cells/mm^3^ (N = 4, n = 7 slices), BA = 21,050 S100β+ cells/mm^3^ (N = 4, n = 6 slices), p = 0.4452). **(E)** Representative maximum intensity projection. Scale bar: 10 μm. **(F, G)** Perimeter (Mann-Whitney test: LA = 239.2 μm (N = 5, n = 16 cells), BA = 262.0 μm (N = 5, n = 19 cells), p = 0.2711) and area (Mann-Whitney: LA = 1,828 μm^2^ (N = 5, n = 16 cells), BA = 2,113 μm^2^ (N = 5, n = 19 cells), p = 0.1332). **(H, I)** Feret diameter definition and quantification (Mann-Whitney test: LA = 64.24 μm (N = 5, n = 16 cells), BA =67.09 μm (N = 5, n = 19 cells), p = 0.2711). Scale bar: 10 μm. **(J)** Representative surface model. Scale bar: 10 μm. **(K-N)** Surface area (Welch’s t test: LA = 134,844 μm^2^ (N = 5, n = 9 cells), BA = 136,640 μm^2^ (N = 5, n = 10 cells), p = 0.9382), volume (Mann-Whitney test: LA = 14,900 μm^3^ (N = 5, n = 9 cells), BA = 27,450 μm^3^ (N = 5, n = 10 cells), p = 0.0340), surface-area-to-volume ratio (Mann-Whitney test: LA = 8.750 μm^-1^ (N = 5, n = 9 cells), BA = 5.620 μm^-1^ (N = 5, n = 10 cells), p = 0.0350) and soma diameter (unpaired t test: LA = 9.118 μm (N = 5, n = 9 cells), BA = 8.441 μm (N = 5, n = 10 cells), p = 0.2824). **(O)** Representative filament model. Scale bar: 10 μm. **(P, Q)** Total process length (unpaired t test: LA = 11,198 μm (N = 5, n = 9 cells), BA = 12,499 μm (N = 5, n = 10 cells), p = 0.4525) and branch-point number (unpaired t test: LA = 3,028 (N = 5, n = 9 cells), BA = 3,468 (N = 5, n = 10 cells), p = 0.3592). **(R)** Three-dimensional Sholl analysis schematic. **(S)** Maximum number of intersections. (Mann-Whitney test: LA = 391.0 (N = 5, n = 9 cells), BA = 444.0 (N = 5, n = 10 cells), p = 0.3154) and **(T)** Sholl profile. **(U)**Heatmap of z-scored group means across cell density and the displayed morphological endpoints. BA volume was greater and surface-area-to-volume ratio lower than in LA; other displayed comparisons were not significant. Bars show group means with SEM and points show individual observations. *p < 0.05; ns, not significant.

### Early-life stress expands BLA astrocyte territory in female mice

We next examined whether early-life stress was associated with altered astrocyte morphology in BLA. In general, stress results in morphological changes across species and brain regions [8,17,19,20,29–37]. While our previous work revealed changes in GFAP density and calcium activity [16], whether early-life stress influences discrete aspects of amygdala astrocyte morphology remain unresolved. To further investigate this question, pups were separated 4 hours a day from their mother with limited bedding from postnatal day 10 to postnatal day 17, where they returned to standard housing conditions. Two weeks later, systemic viral labeling was performed, and a delay of three weeks was respected before morphological analysis (**Fig. 4A-C**). In agreement with our previous findings, astrocyte density with S100β-positive cell density in BLA was unaffected by stress (**Fig. 4D**). In contrast, perimeter and two-dimensional area were increased in stressed females, while

**Figure 4.**
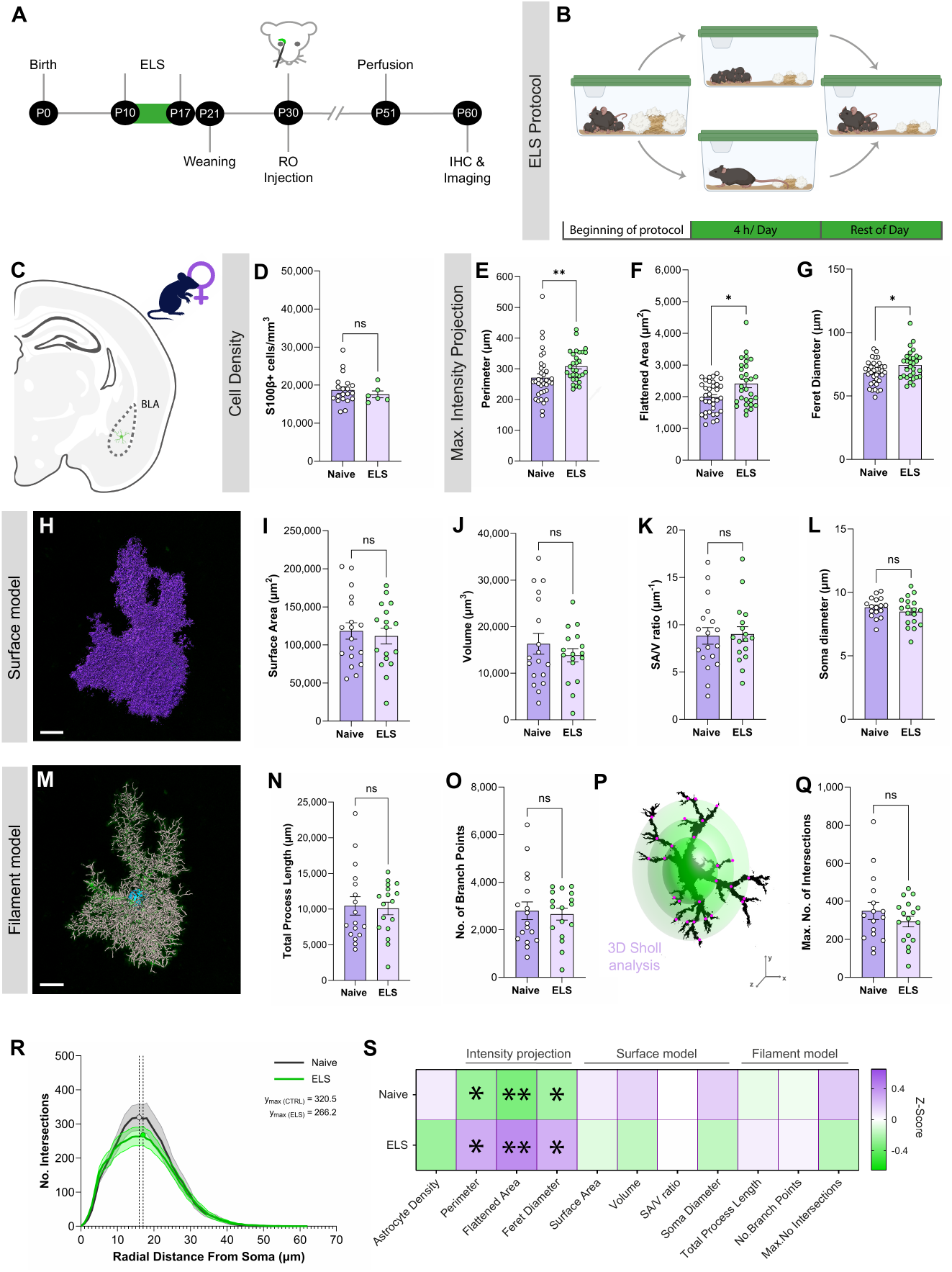
Early-life stress expands BLA territory in female mice. **(A)** Experimental timeline showing early-life stress from P10 to P17, weaning at P21, retro-orbital vector at P30, perfusion at P51 and immunohistochemistry and imaging by P60. **(B)** Maternal separation was applied for 4 h per day, while reduction of bedding lasted 7 days. Initial conditions were restored at P17. **(C)** Female BLA sampling location. **(D)** S100β-positive cell density in naive and stressed female mice (Mann-Whitney test: Naive = 17,800 S100β+ cells/mm^3^ (N = 7, n = 20 slices), ELS = 16,850 S100β+ cells/mm^3^ (N = 3, n = 6 slices), p = 0.4845). **(E-G)** Perimeter (Mann-Whitney test: Naive = 258.0 μm (N = 5, n = 35 cells), ELS = 292.0 μm (N = 5, n = 28 cells), p = 0.0068), area (Mann-Whitney: Naive = 1,957 μm^2^ (N = 5, n = 35 cells), ELS = 2,217 μm^2^ (N = 5, n = 28 cells), p = 0.0436) and Feret diameter (unpaired t test: Naive = 68.32 μm (N = 5, n = 35 cells), ELS = 73.03 μm (N = 5, n = 28 cells), p = 0.0531). **(H)** Representative surface model. Scale bar: 10 μm. **(I-L)** Surface area (unpaired t test: Naive = 118,417 μm^2^ (N = 5, n = 18 cells), ELS = 111,629 μm^2^ (N = 5, n = 17 cells), p = 0.6551), volume (unpaired t test: Naive = 16,320 μm^3^ (N = 5, n = 18 cells), ELS = 13,852 μm^3^ (N = 5, n = 17 cells), p = 0.3635), surface-area-to-volume ratio (unpaired t test: Naive = 8.823 μm^-1^ (N = 5, n = 18 cells), ELS = 9.004 μm^-1^ (N = 5, n = 17 cells), p = 0.8791) and soma diameter (unpaired t test: Naive = 8.823 μm (N = 5, n = 17 cells), ELS = 8.492 μm (N = 5, n = 18 cells), p = 0.3259). **(M)** Representative filament model. Scale bar: 10 μm. **(N, O)** Total process length (unpaired t test: 10,453 μm (N = 5, n = 17 cells), ELS = 10,091 μm (N= 5, n = 17 cells), p = 0.8186) and branch-point number (unpaired t test: Naive = 2,799 (N = 5, n = 17 cells), ELS = 2,660 (N = 5, n = 17 cells), p = 0.7596). **(P)** Three-dimensional Sholl analysis schematic. **(Q)** Maximum number of intersections (Mann-Whitney test: Naive = 320.5 (N = 5, n = 16 cells), ELS = 309.0 (N = 5, n = 17 cells), p = 0.5752). **(R)** Sholl profile. **(S)** Heatmap of z-scored group means across cell density and the displayed morphological endpoints. Perimeter, flattened area and Feret diameter were greater following stress; other displayed summary measures were not significant. Bars show group means with SEM, and points show individual observations. *p < 0.05; **p < 0.01; ns, not significant.

Feret diameter remained similar in both groups (**Fig. 4E-G**). These territorial measures were not accompanied by significant differences in three-dimensional surface area, volume, surface-area-to-volume ratio (**Fig. 4H-K**), or soma diameter (**Fig. 4K**). Next, we examined astrocyte ramification and branching with total process length, branch-point number and maximum Sholl intersections were also unchanged, demonstrating little impact of ELS on astrocyte arborization (**Fig. 4M-Q**). The Sholl curve showed a trend towards a lower peak following stress (**Fig. 4R**). The integrated heatmap summarizes the selective positive shift in two-dimensional territorial data compared with three-dimensional unvarying surface, soma, and filament data in ELS female mice (**Fig. 4S**). These results reveal differences relating to cell territory measurements, whether they were obtained using two-or three-dimensional analysis, with the former demonstrating expanded projected territory following ELS. Furthermore, process complexity remained overall unaffected by the stressor.

### Early-life stress reduces BLA astrocyte size in males

Finally, we investigated the impact of ELS in male mice. Taking animals from the same early-life-stress schedule and applying the same workflow (**Fig. 5A-C**). Once again, S100β-positive cell density did not differ between naive and stressed males (**Fig. 5D**). In contrast to our data from female mice, males showed no significant change in perimeter, two-dimensional area, and Feret diameter (**Fig. 5E-G**). Stress was instead associated with lower three-dimensional surface area and volume (**Fig. 5H-J**). Considering parallel changes in surface area and volume, the surface-area-to-volume ratio was conserved (**Fig. 5K**) and soma diameter was not significantly different (**Fig. 5L**). Similar to female mice, total process length, branch-point number and maximum Sholl intersections were also unchanged (**Fig. 5M-R**). The integrated heatmap emphasizes the negative surface and volume profile relative to the other morphological measurements (**Fig. 5U**). Thus, the male response is characterized by reduced three-dimensional size rather than projected territorial expansion with no detectable change in arbor complexity. This, combined with an unaltered astrocyte density in the BLA, points to a decrease in each astrocyte’s agency over the extracellular space.

**Figure 5.**
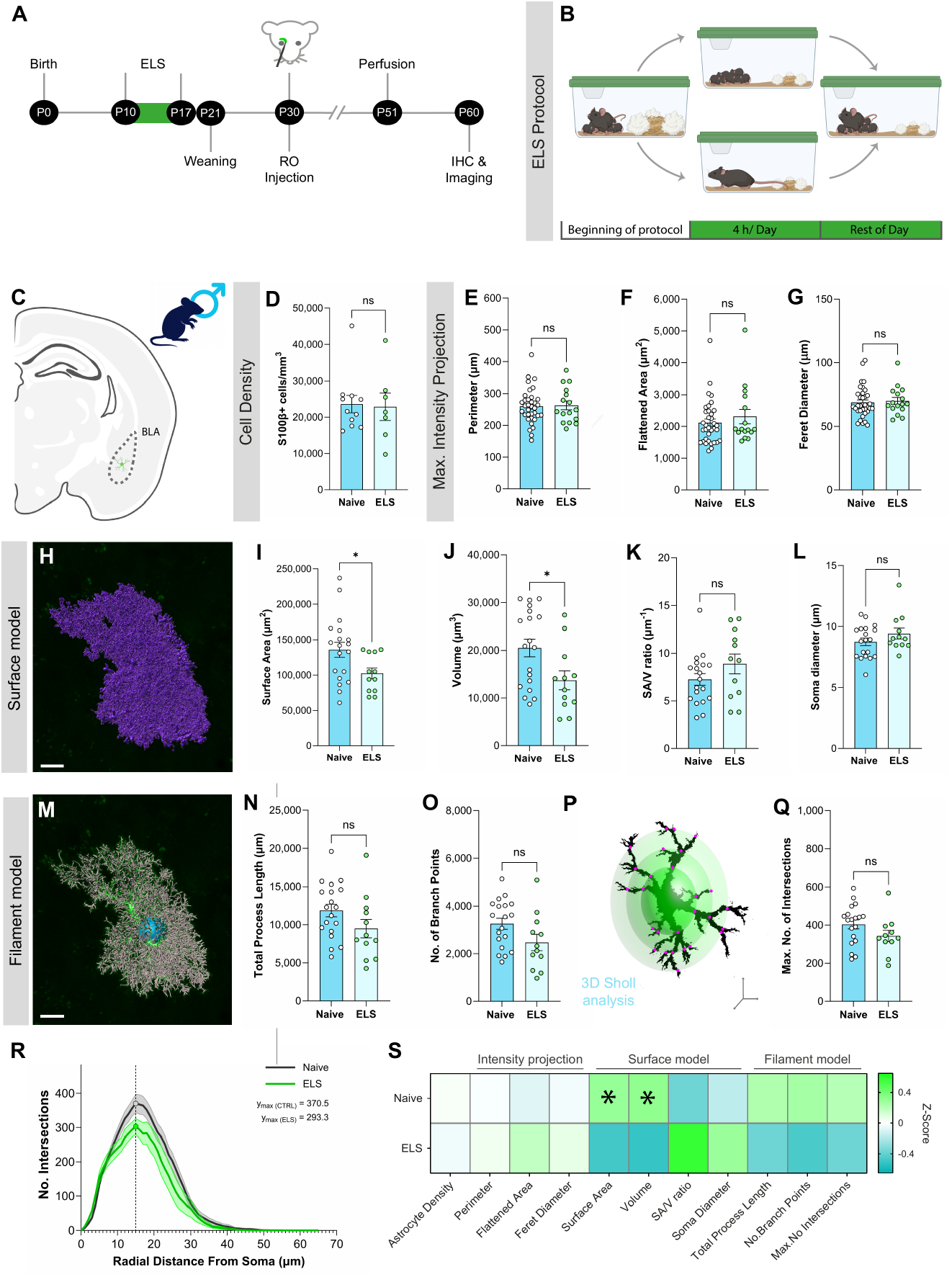
Early-life stress reduces BLA astrocyte size in males. **(A)** Experimental timeline. **(B)** Early-life-stress procedure applied for 4 h per day from P10 to P17. **(C)** Male BLA sampling location. **(D)** S100β-positive cell density in naive and stressed males (Mann-Whitney test: Naive = 21,600 S100β+ cells/mm^3^ (N = 4, n = 13 slices), ELS = 21,400 S100β+ cells/mm^3^ (N = 4, n = 7 slices), p = 0.9385). **(E-G)** Perimeter (unpaired t test: Naive = 260.1 μm (N = 5, n = 35 cells), ELS = 263.1 μm (N = 3, n = 16 cells), p = 0.8558), area (Mann-Whitney test: Naive = 1,958 μm^2^ (N = 5, n = 35 cells), ELS = 1,914 μm^2^ (N = 3, n = 16 cells), p = 0.5531) and Feret diameter (Mann-Whitney test: Naive = 66.27 μm (N = 5, n = 35 cells), ELS = 67.66 μm (N = 3, n = 16 cells), p = 0.4877). **(H)** Representative surface model. Scale bar: 10 μm. **(I-L)** Surface area (unpaired t test: Naive = 135,789 μm^2^ (N = 5, n = 19 cells), ELS = 102,417 μm^2^ (N = 3, n = 12 cells), p = 0.0311), volume (Mann-Whitney: Naive = 20,900 μm^3^ (N = 5, n = 19 cells), ELS = 12,900 μm^3^ (N = 3, n = 12 cells), p = 0.0250), surface-area-to-volume ratio (unpaired t test: Naive = 7.259 μm^-1^ (N = 5, n = 19 cells), ELS = 8.886 μm^-1^ (N = 3, n = 12 cells), p = 0.1591) and soma diameter (unpaired t test: Naive = 8.762 μm (N = 5, n = 19 cells), ELS = 9.433 μm (N = 3, n = 12 cells), p = 0.2063). **(M)** Representative filament model. Scale bar: 10 μm. **(N, O)** Total process length (unpaired t test: Naive = 11,883 μm (N = 5, n = 19 cells), ELS = 9,500 μm (N = 3, n = 12 cells), p = 0.1041) and branch-point number (unpaired t test: Naive = 3,260 (N = 5, n = 19 cells), ELS = 2,470 (N = 3, n = 12 cells), p = 0.0573). **(P)**Three-dimensional Sholl analysis schematic. **(Q)**Maximum number of intersections (unpaired t test: Naive = 403.9 (N = 5, n = 19 cells), ELS = 343.1 (N= 3, n = 11 cells), p = 0.1289). **(R)**Sholl profile. **(U)** Heatmap of z-scored group means across cell density and the displayed morphological endpoints. Surface area and volume were lower following stress; other displayed summary measures were not significant. Bars show group means with SEM, and points show individual observations. *p < 0.05; ns, not significant.

## DISCUSSION

Our previous study examined cell functional characteristics of amygdala astrocytes [13,16], leaving morphological characteristics unexplored. For this purpose, this study uses systemic sparse eGFP labeling to compare astrocyte morphology across amygdala subdivisions, sex, and early-life experience.

Three findings define the current dataset. First, retro-orbital AAV.PHP.eB-GfaABC1D-eGFP provides broad anatomical access while retaining isolated cells that can be analyzed using complementary surface, filament and Sholl workflows. Second, baseline LA-BA differences are sex dependent, with female morphology relatively homogenous between regions, whereas male astrocytes exhibited greater reconstructed volume and lower surface-area-to-volume ratio in the BA compared to LA. Third, early-life stress is associated with different structural responses in female and male BLA astrocytes. Females show greater projected territorial dimensions, whereas males show reduced three-dimensional surface area and volume.

While we used this sparse labelling method to reveal discrete morphological differences in astrocytes across the BLA in control and stress conditions, previous work has also leveraged this approach to sparsely label hippocampal astrocytes for three-dimensional reconstruction [38]. The major practical advantage is anatomical reach. A single injection into the blood-stream produced reporter-positive profiles in cortex, hippocampus, hypothalamus and amygdala, permitting later selection of regions without a separate stereotaxic injection for each target. This approach can be valuable when regional heterogeneity is a primary focus of a research question [4,5,39]. The example here is identification of sex-specific effects of stress on astrocyte structure.

The regional comparisons in non-stressed mice suggest that LA and BA astrocytes are not a homogenous population, but they also show that the regional distinction is sex specific. We found that astrocytes in male mice were larger in the basal compared to lateral amygdala, while no differences were observed between these regions in female mice. While we only assessed morphology, regional variation in astrocyte molecular identity and morphology is well established across the central nervous system [3,4] and is likely to participate in this morphological heterogeneity in basal conditions as well as following stress. The stress experiment revealed a second dissociation. Female BLA astrocytes showed greater perimeter and flattened area without significant increases in surface area or volume. This combination suggests expansion or elongation of the projected envelope rather than uniform three-dimensional hypertrophy. Male BLA astrocytes showed an opposite response consisting of decreased surface area and volume, while projected dimensions and resolved filament measures remained unchanged. Our observations regarding high surface-area-to-volume ratio of astrocytes align with data from ultrastructural work [3,28–30]. Biophysical modelling suggests that this ratio can have physiological relevance as it can impact transmembrane ion flux [31], as such regional differences and the impact of stress could have functional implications for astrocyte-neuron interactions.

The approach also has clear boundaries. Regional expression was heterogeneous, so systemic delivery should not be assumed to yield a uniform labeling density. Certain brain regions, like the cortex, seemed more susceptible to astrocyte labelling, with more sparsity the hippocampus. Inter-individual variability may also play a factor. More importantly, morphology and promoter identity do not by themselves establish complete astrocyte specificity [12]. The representative GFAP image supports an astrocytic interpretation of the labeled profiles, and the reconstructed cells have the expected territorial morphology. These measures will determine how broadly the protocol can be transferred across doses, strains and experimental states. Additionally, because confocal reconstruction does not resolve the full nanoarchitecture of the smallest astrocyte processes, higher-resolution imaging and explicit analysis of reporter intensity and segmentation sensitivity will be needed to distinguish the extent of astrocyte process remodelling, in particular around synaptic elements [3,40,41].

In summary, systemic labeling provides a practical route to spatially distributed single-cell astrocyte morphometry. The current study identifies a selective male regional phenotype under naive conditions and a sex-dependent dissociation in the morphology associated with early-life stress. Females show expansion of projected BLA territory, whereas males show contraction of three-dimensional size. These results indicate that planar, volumetric, and skeleton-based measurements capture related but non-equivalent aspects of astrocyte structure. They also caution against reducing morphology to one label such as hypertrophy or atrophy. Region-, paradigm- and sex-dependent structural responses have also been described in other stress models [8,17,19,20,36,42], with our work adding to this body of literature and providing a useful roadmap for future studies.

## DATA AND CODE AVAILABILITY

Data will be made available at Borealis data depository upon publication - https://doi.org/10.5683/SP4/9NAXIF

## AUTHOR CONTRIBUTIONS

Conceptualization, C.C.A., A.B., and C.M.-R.; methodology, C.C.A., A.B., and C.M.-R.; investigation, C.C.A., A.B., M.G., K.N., and C.M.-R; formal analysis, C.C.A., A.B.; project administration, S.P. and C.M.-R.; writing – original draft, C.C.A and C.M.-R.; writing – review and editing, C.C.A., A.B., M.G., K.N., and C.M.-R; funding acquisition, C.M.-R.

## ACKNOWLEDGEMENTS

We thank lab members for their input. We thank Dr. Aurélie Cleret-Buhot and Dr. Mame-Kany Diop (CRCHUM Cellular Imaging Core) for microscopy training, Véronique Barrès and Liliane Meunier (CRCHUM Molecular Pathology Core) for imaging the coronal brain shown in Figure 1 and the CRCHUM Animal Facility.

This project was supported by funding from the Fonds de Recherche du Québec – Santé (FRQS; 309889); the CHUM Foundation; Canadian Institutes of Health Research (CIHR) project grant (478629); the Natural Science and Engineering Research Council of Canada (NSERC) Discovery grant (RGPIN-2021-03211); the Brain Canada Future Leaders award (BCFA_2024_00796) from the Canada Brain Research Fund (CBRF), an arrangement between the Government of Canada (through Health Canada), the Brain Canada Foundation, and the Azrieli Foundation to C.M.-R. C.C.A was supported by an NSERC undergraduate research scholarship. M.G. was supported by a doctoral fellowship from the Fonds de Recherche du Québec – Santé. C.M.-R. was supported by a Junior 2 Chercheur-Boursier.

